# Genomic repeat landscape evolution across the teleost fish lineages

**DOI:** 10.1101/2023.03.03.530935

**Authors:** William B Reinar, Ole K Tørresen, Alexander J Nederbragt, Michael Matschiner, Sissel Jentoft, Kjetill S Jakobsen

## Abstract

Repetitive DNA make up a considerable fraction of most eukaryotic genomes. In fish, transposable element (TE) activity have coincided with rapid species diversification. Here, we annotated the repetitive content in 100 genome assemblies, covering the major branches of the diverse lineage of teleost fish. We investigated if TE content correlates with family level net diversification rates and found support for a weak negative correlation. Further, we found that TE content, the degree of parental care and short tandem repeat (STR) content contributed to genome size variability. In contrast to TEs, STR content showed a negative relationship with genome size. STR content did not correlate with TE content, which implies independent evolutionary paths. Last, marine and freshwater fish have large differences in STR content. The most extreme propagation was found in the genomes of codfish species and Atlantic herring. Such a high density of STRs is likely to increase the mutational load, which we propose could be counterbalanced by high fecundity as seen in codfishes and herring.

## Introduction

Repetitive sequences including transposable elements (TEs) and short tandem repeats (STRs) comprise large fractions of most eukaryotic genomes. STRs are repetitive stretches of DNA with unit sizes ranging from 1 to 10 bp, increasing and shrinking in size primarily due to replication slippage (Levinson & Gutman 1987). The origin of STRs in genomes can be attributed to processes of unequal crossing over (Smith 1976). New STRs can also originate from parts of active TEs, as insertions of poly-A tails from retrotransposition, or from *de novo* mutations of STR-like patterns (Ellegren 2004; Pasquesi et al. 2018). TEs take advantage of the DNA replication and transcription processes of their hosts to facilitate propagation and are defined into two main classes: DNA transposons, which transpose directly from DNA to DNA, and retrotransposons (RTs) that transpose *via* an RNA intermediate. RTs are further divided into elements containing long terminal repeats (LTRs) and those that do not, the long interspersed nuclear elements (LINEs) and the short interspersed nuclear elements (SINEs) (Kapitonov & Jurka 2008).

Comparative studies have revealed that TE content to some extent explains genome size variation across vertebrates (Chalopin et al. 2015) and across chordates (Canapa et al. 2015). Within more phylogenetically narrow taxa, differences in repeat content do not necessarily reflect the variation in genome size, such as within reptiles, mammals and birds (Pasquesi et al. 2018; Kapusta et al. 2017). In the largest vertebrate group, teleost fish, the correlation between genome size and repetitive DNA content appears to be modest (Gao et al. 2016; Yuan et al. 2018; Canapa et al. 2015; Chalopin et al. 2015), with the largest study (Yuan et al. 2018) reporting an R of 0.6 (R^2^: 0.36). In contrast, TE content have been suggested to explain 98% of the variation in genome size in angiosperms (Tenaillon et al. 2010). Due to the nature of TE propagation it is not surprising that an increase in TE copies may lead to an increase in genome size, but the empirical evidence is less clear for correlations between STR content and genome size. Across eukaryotic domains, the relationship seems to be positive (Hancock 2002; Mayer et al. 2010; Hancock 1996). However, no significant correlation has been reported within domains (Morgante et al. 2002). As reported by Hardie and Hebert (2004), genome size is likely linked to differences in egg diameter, parental care and aquatic habitat (saltwater or freshwater). These factors have so far not been taken into account when testing the relationship between genome size and repetitive DNA in teleosts.

Beyond their contribution to genome size variability, TEs have been postulated to cause deletions, translocations, duplications, and inversions in response to stress conditions (McClintock 1984). A role for TEs in adaptation has been indicated in invasive species of ants, where TE-dense genomic islands were shown to have generated variability in genes deemed important in the adaptation process (Schrader et al. 2014). Interestingly, bursts of TE activity coinciding with speciation have been found in studies of a variety of taxa (Rebollo et al. 2010), including mammals (Ricci et al. 2018). Within teleost fish, elevated TE activity has been shown to coincide with species radiations in salmonids (de Boer et al. 2007) and cichlids (Salzburger 2018; Brawand et al. 2014). Beside a potential role in adaptive radiations through generating adaptive mutations, a mechanism of which TEs could influence speciation is by causing chromosomal rearrangements, possibly as a response to epigenetic release due to environmental stress (Rebollo et al. 2010), which in turn can lead to reproductive isolation. For STRs, different length variants present in a population contribute to the genetic variation and have been shown in some cases to be functionally relevant (Gemayel et al. 2015; Press et al. 2018; Gymrek et al. 2016). As with TEs, STR content varies across vertebrates, with frequencies from approximately 100 loci/Mbp to 1000 loci/Mbp and densities from 1000 bp/Mbp to 50 000 bp/Mbp (Adams et al. 2016; Tørresen et al. 2017, 2018). A large proportion of these STRs occur outside genes; however, in humans for instance, around 4500 STRs occur in protein coding regions (Willems et al. 2014). A STR within an open reading frame (ORF) often encodes homo- or di-amino acid tracts that to a large extent overlap with intrinsically unstructured protein regions (Simon & Hancock 2009). Such regions are abundant in proteins that interact with other proteins (Huntley & Clark 2007). On the other hand, STRs occurring in regulatory regions can affect the expression of genes (Vinces et al. 2009; Quilez et al. 2016) and STRs in introns may impact RNA splicing (Hefferon et al. 2004; Press et al. 2018).

In light of the above-mentioned observations, a key question is to what extent the genomic repeat landscape impacts the evolution of vertebrates, here exemplified by teleost fishes. First, we investigated the interplay between genome size, aquatic habitat, parental care and repetitive DNA content, using comparative methods taking phylogenetic relationships as well as assembly quality into account. Next, we focused on diversification. Our focal group, teleosts, is the most species rich group of all vertebrates and serves as a suitable system to test for associations between the TE/STR landscape and diversification, given the recent genomic sequencing initiatives of multiple teleost species (Malmstrøm et al. 2016, 2017; Musilova et al. 2019) as well as available species richness data. Teleostean families differ widely in species diversity, ranging from monotypic families such as Helostomatidae to the Cyprinidae, with ∼3000 species. Teleosts display a 10-fold difference in TE content (Gao et al. 2016, 2017; Chalopin et al. 2015; Canapa et al. 2015) and a 13-fold difference in STR content (Tørresen et al. 2018). We annotated the TE and STR content in the genome assemblies of 100 teleost fish and one non-teleost ray-finned fish (spotted gar, *Lepisosteus oculatus*). Our samples cover the major teleost branches, allowing us to describe differences in TE and STR content after ∼270 million years of evolution, and to investigate the role of repetitive DNA in teleost genome size evolution and its potential influence on diversification.

## Results

### Substantial changes in TE content over 270 million years of evolution

Our annotation of TEs (and unclassified interspersed repeats) revealed high variation among species (Figure 1). DNA transposon content varies the most, ranging from 1.6% in the tetraodon (*Tetraodon nigroviridis*) genome to 37.1% in the zebrafish (*Danio rerio*) genome. The LTR-RT content ranges from 0.48% in bluefin trevally, southern platyfish and climbing perch (*Caranx melampygus, Xiphoporus maculatus* and *Anabas testudineus*) to 7.4% in opah (*Lampris guttatus*). LINE content varies from 0.89% in blind cavefish (*Astyanax mexicanus*) to 12.6% in giant oarfish (*Regalecus glesne*) and SINE content ranges from 0.02% in electric eel (*Electrophorus electricus*) to 3.6% in giant oarfish (*R. glesne*). We also quantified the proportions of DNA transposons, LTR retrotransposons, LINEs and SINEs relative to the total classified TE content. We find that DNA transposons collectively make up the largest proportion of the TE composition in most teleost fish genomes (94 out of 101 species). However, we find multiple lineage-specific differences in TE composition. The DNA transposon fraction is especially high in the genomes of *Astyanax mexicanus* (89.7%), Cyprinidae (mean: 77.5%), Sebastidae (mean: 76.9%) and Poeciliidae (mean: 74.1%). Of retrotransposons, LINEs are the most prevalent TE subclass, and display the highest relative fractions in *Cetomimus* sp. (51.4%), *Regalecus glesne* (51.0%), Tetraodontidae (mean of 44.9%) and *Lampris guttatus* (40.9%). The LTR-RT fraction is comparably low in most of the genomes studied, but is more prevalent in *Gasterosteus aculeatus* (25.2%), *L. guttatus* (24.2%) and Gadidae (mean: 23.3%). Relative SINE fractions are generally low (mean: 4.2%), with a few exceptions being *Synodus synodus* (16.8%), the non-teleost *Lepisosteus oculatus* (16.5%) and *R. glesne* (14.5%). The Tetraodontidae family (represented by *Takifugu rubripes, Takifugu flavidus* and *Tetraodon nigroviridis*) have a particularly small fraction of DNA transposons, a feature shared only with distant relatives such as *L. oculatus, R. glesne* and *Cetomimus sp*. The two lampriform fishes (*R. glesne* and *L. guttatus*) stand out from other fishes in TE composition, and from each other as well. Overall, the large differences in TE composition among and sometimes within teleost families highlight the dynamic nature of TEs during teleost evolution.

**Figure 1.**
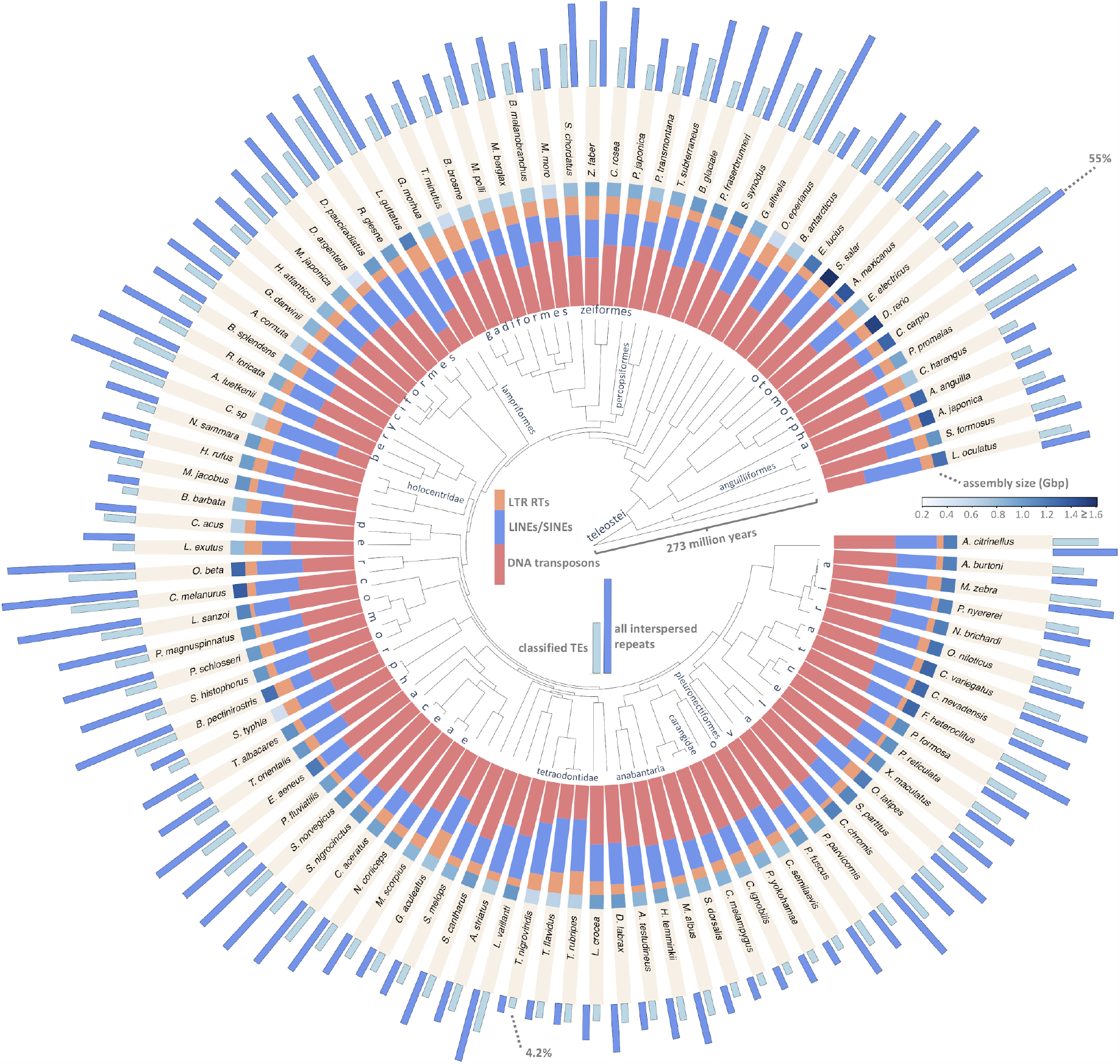
Transposable element (TE) content in 101 fish genomes. The phylogeny is the same as shown in Musilova et al. 2019. Stacked colored bars show the relative proportions of TEs; LTR retrotransposons (orange), LINEs and SINEs (blue) and DNA transposons (red). Boxes between the stacked bars and species names are colored according to assembly size, serving as a proxy for genome size. The light blue and dark blue outer bars show the genomic percentage of classified TEs and all interspersed repeats (unclassified repetitive elements) respectively, with the longest bars (*D. rerio*) representing 48.0% classified TEs or 55% interspersed repeats, and the shortest (*T. nigroviridis*) representing 4.2% classified TEs. Some taxonomic clades are named.

### Interplay between genome size, repetitive DNA and other factors

We performed phylogenetic generalized least square (PGLS) regression to test if genome assembly size was correlated with the TE and STR content of the assemblies, while taking the phylogenetic relationships among samples into account, as well as the aquatic habitat and degree of parental care. The correlation between the number of TEs and genome assembly size (R^2^: 0.72, *p* value: < 2.2e^-16^, Figure 2a) was stronger than between the genomic proportion of TEs and genome assembly size (R^2^: 0.15, *p* value: 3.6e^-5^, Figure 2c), the latter being a more direct measure of the influence on genome size. The number of STRs displayed a positive correlation (R^2^: 0.41, *p* value: 5.9e^-13^, Figure 2b), but the genomic proportion of STRs appeared to have a negative relationship with genome assembly size. This apparent relationship did not reach a significance threshold of 5% for a linear relationship (R^2^: 0.022, *p* value: 0.078, Figure 2d). As the local regression resembled a negative exponential relationship, we log_10_-transformed the STR content of the assembly and repeated the test, yielding a weak linear negative correlation (R^2^: 0.044, *p* value: 0.021). We continued using log_10_-transformed STR content in all other tests. For all tests with genome assembly size as a response, we omitted the Atlantic salmon (*Salmo salar*) and zebrafish (*Danio rerio*), as both drastically impacted the regressions when included (plots including these species can be viewed in Supplementary Figure 1). Although the variation seen in *S. salar* and *D. rerio* is biologically meaningful, *S. salar* is an extreme outlier in terms of genome size (∼ 3 Gb) due to a salmonid-specific whole genome duplication and the *D. rerio* assembly has a N50 of > 1 Mb, while no other assembly reached a N50 of 100 Kb. We did not find any correlation between the genomic proportion of TEs and the genomic proportion of STRs. Next, we modelled genome size as a response to TE content, (log_10_) STR content, aquatic habitat (marine/freshwater), degree of parental care. To control for differences in assembly quality, we included assembly quality metrics as covariates. We used data on aquatic habitat and parental care from FishBase (Froese & Pauly 06/2018), where the degree of parental care is defined according to (Balon 1990). We grouped fish that carries eggs in their mouth or body (bearers) and that guard their eggs in nests or similar (guarders) together. In the full model (Table 1), aside from TE content having a significant positive effect (*p* value: 5.5e^-7^) on genome assembly size and (log_10_) STR content having a significant negative effect (*p* value: 0.02), non-guarding behavior was positively correlated to genome size (*p*: 0.007), while the marine habitat did not have a significant contribution (*p* value: 0.7). The assembly quality metrics N50 and gene completeness had small, but significant effects on the regression. In total, the significant variables explained 47.5 % of the variation in genome assembly size.

**Figure 2.**
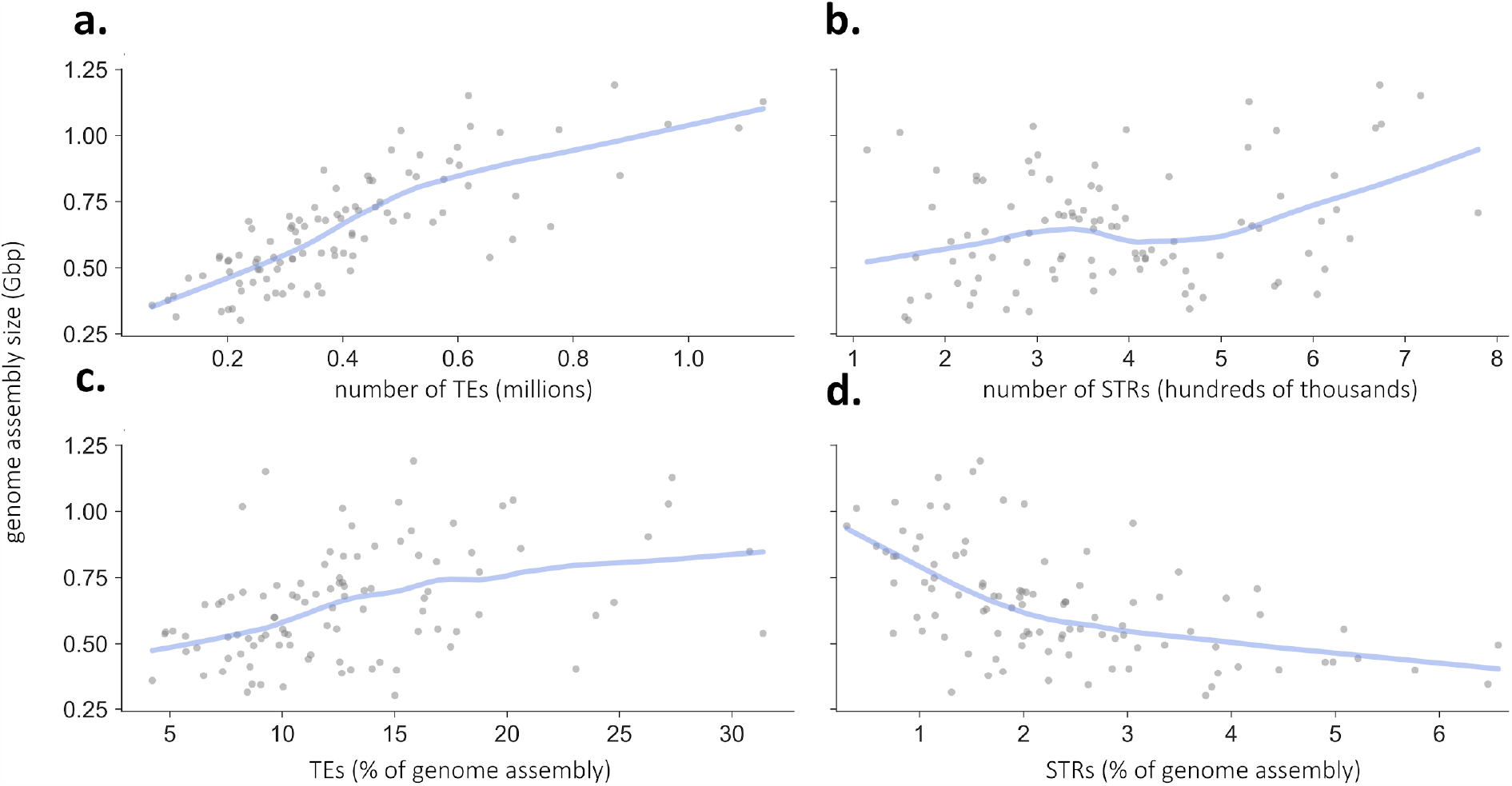
Correlations between repetitive DNA and genome assembly size. The number of classified transposable elements (TEs) (a), the number of short tandem repeats (STRs) (b), the proportion of classified transposable elements (c) and the proportion of STRs (d) covering teleost fish genome assemblies. The light blue lines show the local regression. Plots including Atlantic salmon (*S. salar*) and zebrafish (*D. rerio*) can be seen in Supplementary Figure 1

**Table 1.** Phylogenetic generalized linear model of genome assembly size. Genome assembly size is modelled as a response to the percentage of genomic TE, the (log_10_) genomic percentage of STRs, whether or not the fish species is guarding its eggs, aquatic habitat, and genome assembly metrics (N50 and BUSCO single copy gene completeness).

|  | Explanatory variable | Estimate (Mbp) | Std. error | *t* value | *p* value |
| --- | --- | --- | --- | --- | --- |
| Repetitive elements | percentage TEs | 22.0 | 3.9 | 5.6 | 5.5e<sup>-7</sup> |
|  | log<sub>10</sub> percentage STRs | -238.4 | 102.6 | -2.3 | 0.02 |
| Ecological factors | non-guarding behavior | 115.6 | 41.5 | 2.8 | 0.007 |
|  | marine habitat | -24.4 | 60.5 | -0.4 | 0.7 |
| Assembly quality metrics | N50 contig | -0.003 | 0.001 | -2.5 | 0.01 |
|  | gene completeness | 0.1 | 0.03 | 4.6 | 2.5e<sup>-5</sup> |

#### STR variation across teleost lineages linked to aquatic habitat

Our STR annotation efforts show that there is high variability in STR content within teleost fish, both with respect to total STR content and relative differences of STRs with different unit sizes (Figure 3a). One striking pattern is the proportion of STRs with unit size 5-10 in Gobiidae (Chatrabas melanurus, Lesueurigobius cf. sanzoi, Periophthalmus magnuspinnatus, Periophthalmodon schlosseri, Scartelaos histophorus and Boleophthalmus pectinirostris), more specifically decanucleotide repeats. Suspecting that this might be an artifact, we looked at Gobiidae tandem repeats with unit sizes from 1 to 20, and found that the high proportion of decamers actually represents a high proportion of k-mers with unit sizes 10-20 (mostly 11-mers), which likely confuses the repeat detection algorithm (Phobos) when repeats are interrupted. Why Gobiidae have such a unique STR landscape compared to other teleosts requires further investigation. Using PGLS, we found a significant difference between marine and freshwater fish in STR content (p: 0.0003, Figure 3b), supporting the tendency found in Yuan et al. (2018). The association was robust to removal of the whole Ovelentaria clade, which mainly contain freshwater fish. We noted that families within the codfishes (Gadidae, Lodidae, Merluccidae, Macrouridae, Bathygadidae and Moridae) have particularly high STR content, compared to the other species. By annotating additional codfish assemblies (from Malmstrøm et al. 2016, 2017) we found that extreme STR propagation is common within this lineage (Figure 4).

**Figure 3.**
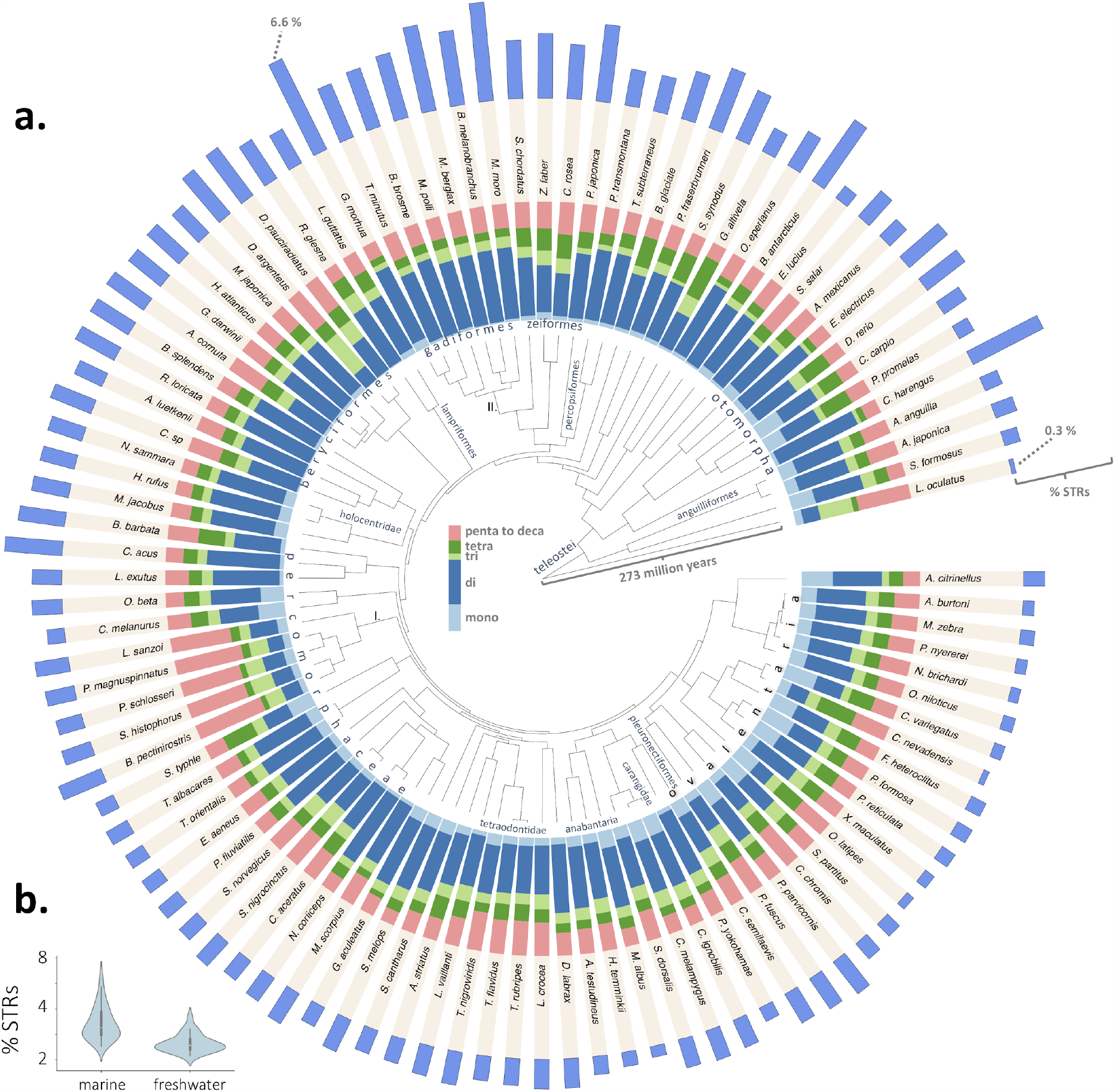
Lineage-specific variation in STRs linked to habitat. (a) Short tandem repeat (STR) content in the genomes of 100 teleost fish and one non-teleost (*L. oculatus*). The phylogeny is the same as in Figures 1. The stacked bars show the relative distribution of STRs grouped by unit size (from mononucleotide repeats with unit size of one to four, and for clarity, grouped data for STRs with unit sizes from five to ten). The outermost bars show total STR content, with the highest bar (Atlantic cod, *Gadus morhua*) representing 6.6% STR content (with the Gadiformes in general having high STR content) and the lowest bar (spotted gar, *Lepisosteus oculatus*) indicating 0.3% STR content. Gobiidae (I.) and Gadiformes (II.) are highlighted as they are mentioned in the main text. (b) Violinplots showing the difference in STR content between marine and freshwater genomes.

**Figure 4.**
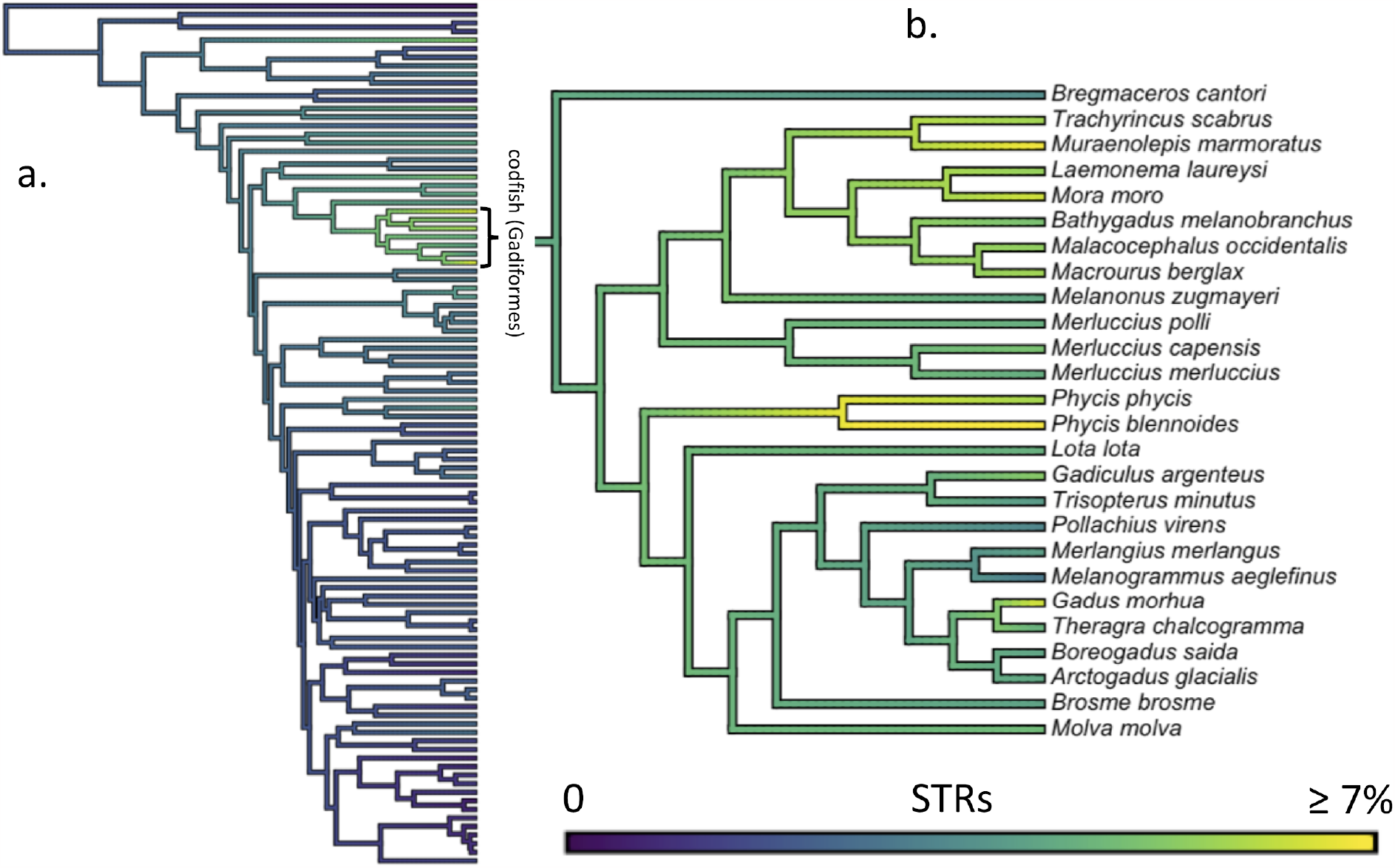
STR content in the codfish lineage. (a) The 101 species phylogeny from Musilova et al. 2019, with the codfish (Gadiformes) highlighted. Branches are colored by maximum likelihood ancestral state estimates of genomic STR percentage. (b) Additional codfish species depicting the phylogeny of Gadiformes from Malmstrøm et al. (2016). Extreme STR values are found within this order, exceeding 9.5% in greater forkbeard (*Phycis blennoides*). For clarity, the maximum cutoff was set to 7%.

### TE content not positively correlated with net diversification

To test if repetitive DNA content within the genomes of teleost families are associated with net diversification rates, we used estimates from Scholl and Wiens (2016) where family-specific net diversification rates were calculated across the tree of life, including 45 out of 71 of our surveyed teleost families (see Material and Methods). Family-specific net diversification estimates were regressed on median TE and STR content per family, as well as median genome assembly size. We used median instead of mean TE proportions per family to avoid that extreme outliers, such as the zebrafish, distort the numbers for individual families. PGLS regressions showed that genome assembly size does not correlate with diversification (Table 2, Figure 5a). We found no significant correlation between net diversification and STR content (Table 2, Figure 5b). For TEs, we did not find support for a positive correlation, but rather a weak negative correlation between net diversification rates and both only classified TE content (R^2^: 0.07, *p* value: 0.015, Figure 5c). The correlation remained significant after removing Pleuronectidae, which in our dataset is an outlier in terms of net diversification rate (classified TE content: R^2^: 0.07, *p* value: 0.019). Tests were repeated using net diversification rates based on different assumed extinction rates (0.1, 0.5 and 0.9), and with total amount of interspersed DNA instead of the percentage of classified TEs, which had negligible impacts on the results.

**Table 2.**
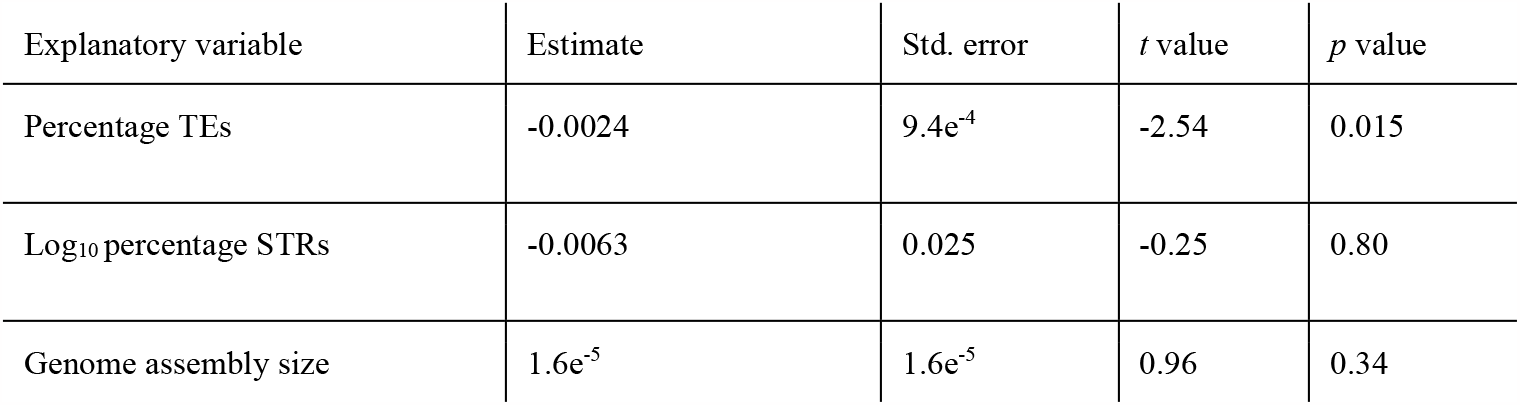
Phylogenetic generalized linear model of family-specific net diversification rates with an assumed extinction rate of 0.5. The median net diversification rate per family is modelled as a response to the median family-specific genomic percentage of TEs, the (log_10_) genomic proportion of STRs and genome assembly size.

**Figure 5.**
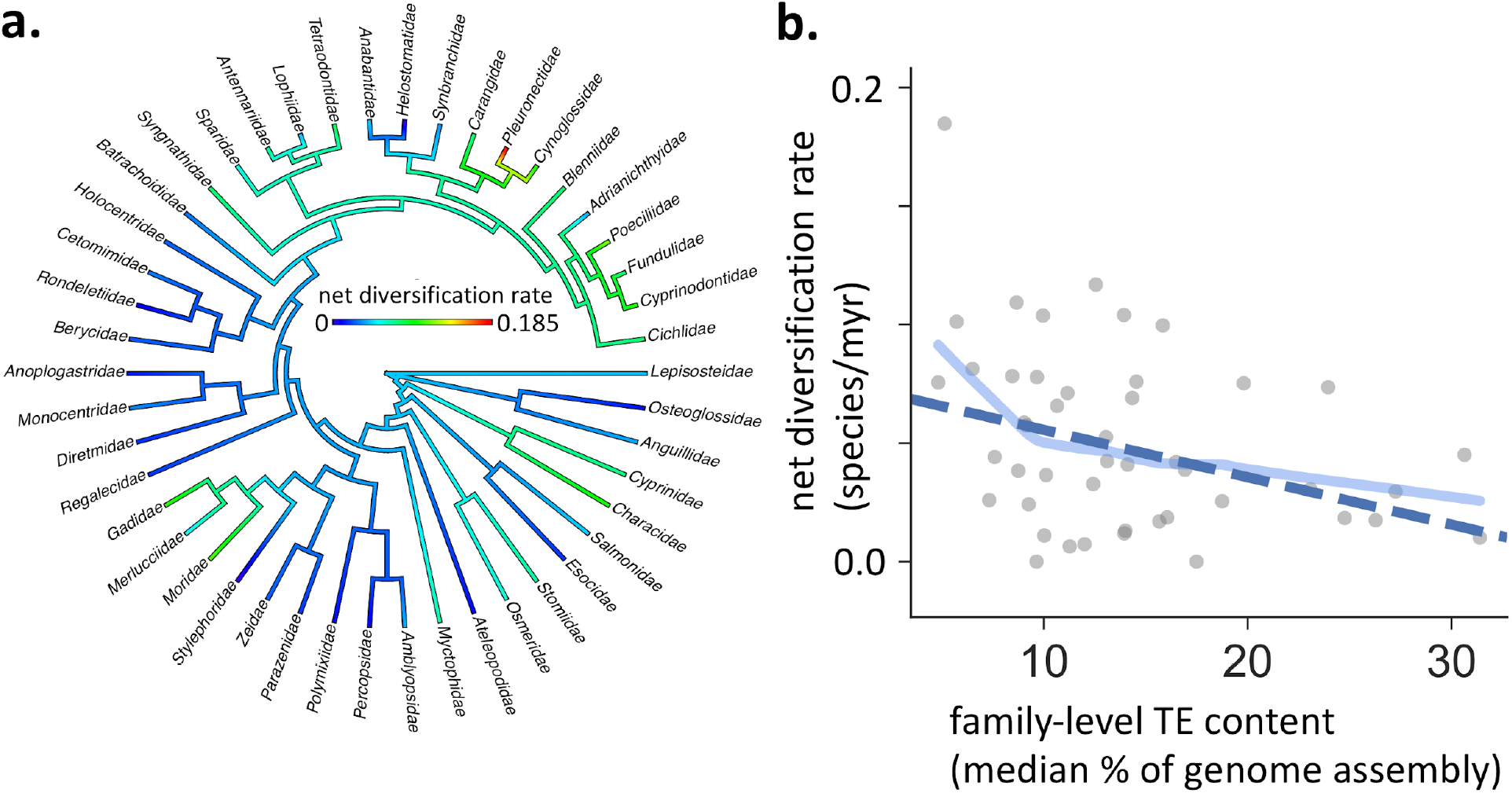
Net diversification rates of teleost fish families (n = 45) do not show a positive correlation with median TE content. (a) Family-specific net diversification rates as estimated by Scholl and Wiens 2016. (b) Phylogenetic generalized between median TE content and median net diversification rate. (d) The light blue lines show the local regression and blue dashed lines show the PGLS regressions.

## Discussion

Using a time-calibrated phylogeny we have investigated the genomic repeat landscape across the teleost radiation. Overall TE content was not positively associated with net diversification, but significantly contributes to genome size variation. Genome size variation was also explained by the genomic percentage of STRs and the degree of parental care. High STR content was associated with smaller genomes, marine habitat and high fecundity (such as codfish and Atlantic herring). We do not observe the same patterns of STR and TE propagation in teleost lineages, pointing towards independent evolutionary paths for these types of repeats.

Our results on the contribution of TEs to genome size variation (Figure 3a-c) support the general tendency observed in chordates (Canapa et al. 2015), vertebrates (Chalopin et al. 2015) and previous studies of teleosts (Yuan et al. 2018). There was, however, large variance, resulting in a fairly low R^2^ of 0.15 (for classified TEs only) and 0.25 (for all interspersed repeats, Supplementary Figure 3). This shows that in teleosts, differential abundance of TEs alone might explain 15% to 25% of the variance in genome size, when the phylogenetic relationship between samples is taken into account. Our model that included STR content, genome assembly quality metrics, and also parental care, which previously have been linked to genome size differences in teleosts (Hardie & Hebert 2004), explained 47.5% of the genome size differences in our samples. Given that the extent of parental care and egg size are positively correlated (Kolm & Ahnesjo 2005), and egg size is positively correlated with genome size (Hardie & Hebert 2004), we expected to find a positive correlation between parental care and genome size. Contrary to expectation, non-guarding behavior had an overall positive contribution to genome size variability. Further, a marine environment did not explain any difference in genome size when the phylogenetic relationship was taken into account.

In comparison, STR content was significantly higher in marine fish (Figure 4), with the most extreme being the codfish (Figure 5). Given the current understanding of STRs as hypervariable regions with occasional functional impact, we speculate that marine species with high fecundity and high mortality of eggs (Duarte & Alcaraz 1989), more robustly tolerate the mutational load of STRs, which is likely substantial. Theory predicts (Nei 2013; Graur 2017) that the number of offspring an individual on average needs to produce to keep the population size constant is a function of the deleterious mutation rate and the number of functional mutable sites. It is likely that STRs increase the deleterious mutation rate and that the extent would depend on the STR mutation rate and the fraction of STRs in functional regions. This could serve as an explanation for why we see elevated STR propagation in marine clades, i.e., fish with more numerous eggs, compared with freshwater fish. In particular, of species with available fecundity estimates (scaled for body size), *G. morhua* and *C. harengus* have the highest fecundity in our dataset (Barneche et al. 2018) and also stand out as having high STR content.

We show that DNA transposons are the most common TEs in teleosts (Figure 2), confirming the pattern observed in other studies. Overall, variation is high across lineages and indicates substantial TE activity over 270 million years of evolution. As elevated TE activity has been shown to coincide with teleost species radiations, such as in salmonids (de Boer et al. 2007) and cichlids (Salzburger 2018; Brawand et al. 2014), and in light of the ongoing discussion of the role of TEs in evolution (Brunet & Doolittle 2015; Doolittle & Brunet 2017; Doolittle & Sapienza 1980), a main objective of this study was to investigate if clades with high TE content have had comparably high net diversification. The test relies on the assumption that a fish family with a high proportion of repetitive elements in their genomes is likely to have had more propagation of repetitive elements than a fish family with a low proportion. Our results do not support that high TE content is linked to higher net diversification rates, but rather show mild support for the contrary (Figure 4a), and we see no apparent pattern with regard to the effect of genomic STR proportions or genome size, at least across our broad selection of teleostean families. This does not rule out that TE insertions can lead to novel adaptive traits, and might facilitate diversification in certain teleost clades, as indicated in studies of African cichlids (Brawand et al. 2014; Santos et al. 2014; Salzburger 2018). However, a general speciation promoting role for TEs is not reflected in our results.

Throughout the study, we assessed TE content to be the sum of interspersed repetitive elements judged by our tools (BLAST, BLASTX and HMMR) to be a TE. The classification process is limited by the extent of prior annotated TEs, which in teleosts are biased towards *D. rerio*. This is illustrated by the values obtained from zebrafish, that has the most extensive prior annotation, and the percentage of classified TEs is (48.0 %) is very close to the total interspersed repeats (52.2 %), which is not the case for most other surveyed fish (see Figure 1). The total amount of repetitive elements includes all classified TEs, non-classified TEs, and some sequences that may not be TEs (but occur in multiple loci). Including this measure, as we did in the test with diversification rates, could serve as a less biased estimate of total TE content. Either way, the amount of detected sequences is strongly correlated with the amount of classified sequences (PGLS R^2^: 0.69, Supplementary Figure 2). The detection of interspersed repeats is not biased by *a priori* information, but can be influenced by assembly quality (Treangen & Salzberg 2011; Simpson & Pop 2015). However, in our models that include multiple covariates, we found that two common assembly quality metrics; contig N50 and gene completeness did not impact our conclusions. It should further be noted that genomes inhabiting high numbers of identical TEs (i.e. families that recently expanded) are expected to be harder to assemble, as identical sequences create collapsed repeats. This can lead to an underreporting of elements in genomes with recent expansions. It is also known that high STR content in combination with short read sequencing can produce assemblies of lower quality, reported in the sequencing efforts of the Atlantic cod (*G. morhua*) genome (Tørresen et al. 2017). This implies that assemblies with low assembly quality likely are underestimated with regards to STR content.

Regardless of some limitations, our results suggest that high proportions of TEs are not positively correlated with net diversification rates in teleost clades, and that elevated levels of STRs are linked to and must thus be tolerated by marine teleosts, potentially due to higher fecundity. Such a link would be very important for understanding genome evolution, but needs to be further investigated within teleosts, as well as in other organism groups.

## Material and Methods

### Genome assemblies and phylogenies

Most genome assemblies were retrieved from a recent teleost genome data release (Malmstrøm et al. 2017), and additional assemblies were sequenced and assembled by Musilova et al. 2019, which also released the 101-species phylogeny. Some genome assemblies were retrieved from ENSEMBL and NCBI. For an overview of assembly origins, see Musilova et al. 2019 and Supplementary Table 1. The codfish phylogeny was taken from Malmstrøm et al. 2016. Details regarding the phylogeny construction can be found in these respective studies.

### TE and STR annotation

For TE annotation, we used a variant of the computational pipeline that is more thoroughly described in (Tørresen et al. 2017), available at https://github.com/uio-cels/Repeats. The pipeline includes multiple TE detection steps using different tools, steps for removing non-TEs from the detected sequences and steps for classifying the elements. For the initial detection step, we used RepeatModeler (v. 1.0.8) (Smit & Hubley 2008-2015) and LTRharvest (part of GenomeTools v. 1.5.7) (Ellinghaus et al. 2008).

RepeatModeler detects all sorts of repetitive sequences and LTRharvest is specialized for detecting LTR-RTs. Using BLASTX, TEs with sequences matching known non-TEs in UniProtKB/Swiss-Prot were removed. To classify the TEs, we used RepeatClassifier, which is a part of the RepeatModeler software. As the tool did not manage to classify all of the remaining sequences, additional similarity searches were performed between the sequences and a curated library of TE sequences (RepBase v. 20150807), using nucleotide BLAST. Finally, we built Hidden Markov Model profiles from the detected sequences using HMMER (v. 3.1b1) (Wheeler & Eddy 2013) and compared the profiles with HMM profiles from databases downloaded from GyDB.org (Llorens et al. 2011) and dfam.org (Hubley et al. 2016), using the nhmmer feature included in HMMER. This resulted in additional sequences being classified at the class and subclass level. We merged the *de novo* TE library with a library of known eukaryotic TEs (RepBase) and used this as input for RepeatMasker (v. 4.0.6), run with the -s (sensitive) option. The .out and .tbl files produced by RepeatMasker served as the basis for the downstream analysis, performed using custom Python scripts. For detection of short tandem repeats we used Phobos v3.3.12 (Mayer et al. 2010) to detect all STRs with unit size 1–10 bp in the genome assemblies. The output was in Phobos native format which was further processed with the sat-stat v1.3.12 program, yielding files with statistics and a GFF file. Other options were set as in Tørresen et al. (2017). For the gobiidae genomes, we ran Phobos with unit sizes 1-20 bp.

### Diversification rates

We retrieved estimates of net diversification rates from Scholl and Wiens (Scholl & Wiens 2016), who calculated diversification rates based on the stem ages of teleost families from the teleost phylogenetic tree produced by Betancur-R et al. (Betancur-R. et al. 2013). They used the method-of-moments estimator as described by Magallon and Sanderson (Magallón & Sanderson 2001);

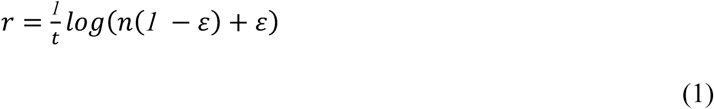

where *r* is the net diversification rate estimate, *t* is the family stem age, *n* is the number of extant species and *ε* is the relative extinction rate. *ε*is included to correct for unsampled, extinct clades. The estimates used in this study are based on the *r* values when *ε* was set to 0.1, 0.5 and 0.9. Note that more recent diversification estimates are available (Rabosky et al. 2018), but cover only marine fish.

### Comparative phylogenetic analyses

Statistical analysis was performed using phylogenetic least-squares (PGLS) regressions using the R package ‘caper’ v. 1.1.0 (Orme et al. 2012). PGLS is a commonly used method for incorporating phylogenetic information in the modelling of associations between traits. PGLS assumes that more closely related species have more similar traits and uses the expected covariance structure to modify the slope and intercept estimates. For tests with net diversification rates we used a pruned phylogeny containing tips representing teleost family stem ages, and used median values per family for all covariates. In all tests, we optimized branch length transformations using maximum likelihood. LOWESS (locally weighted linear regressions) lines were created using the ‘seaborn’ Python package with the ‘regplot’ function and standard parameters.

## Code and data availability

Summaries of the annotation of TEs and STRs, along with the ecological data, are in Supplementary Table 1. Species-specific annotations of TEs and TE-derived DNA can be found at: https://doi.org/10.6084/m9.figshare.8280800 (∼4.6 Gb). The R script used for statistical analysis, can be found at https://github.com/uio-cels/teleost-repeats.

## Acknowledgments

The authors would like to thank Jostein Starrfelt, Masahito Tsuboi and Kjetil Lysne Voje (CEES, University of Oslo) for conceptual input regarding diversification rates. All computational work was performed on the Abel Supercomputing Cluster (Norwegian metacenter for High Performance Computing (NOTUR) and the University of Oslo) operated by the Research Computing Services group at USIT, the University of Oslo IT-department (http://www.hpc.uio.no/). Sequencing library creation and high throughput sequencing was carried out at the Norwegian Sequencing Centre (NSC), University of Oslo, Norway. This research was supported by the Norwegian Research Council under the projects “Functional and comparative immunology of a teleosts world without MHC II (#222378)” and “Evolutionary and functional importance of simple repeats in the genome (#251076)” both led by KSJ. We have adhered to all local, national and international regulations and conventions, and we respected normal scientific ethical practices.

## Supplementary Figures

**Supplementary Figure 1.**
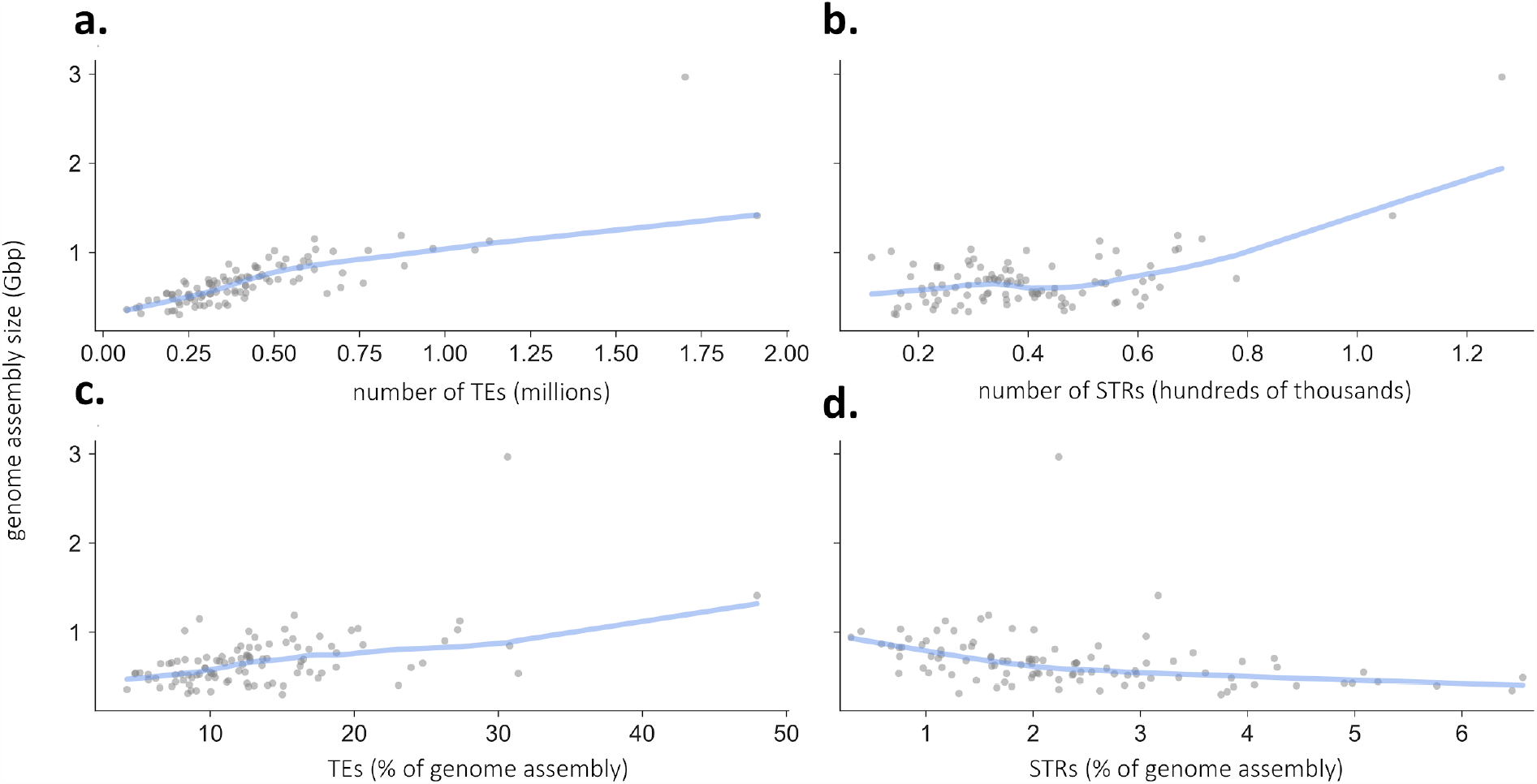
As Figure 1, with Atlantic salmon (*S. salar*) and zebrafish (*D. rerio*) included.

**Supplementary Figure 2.**
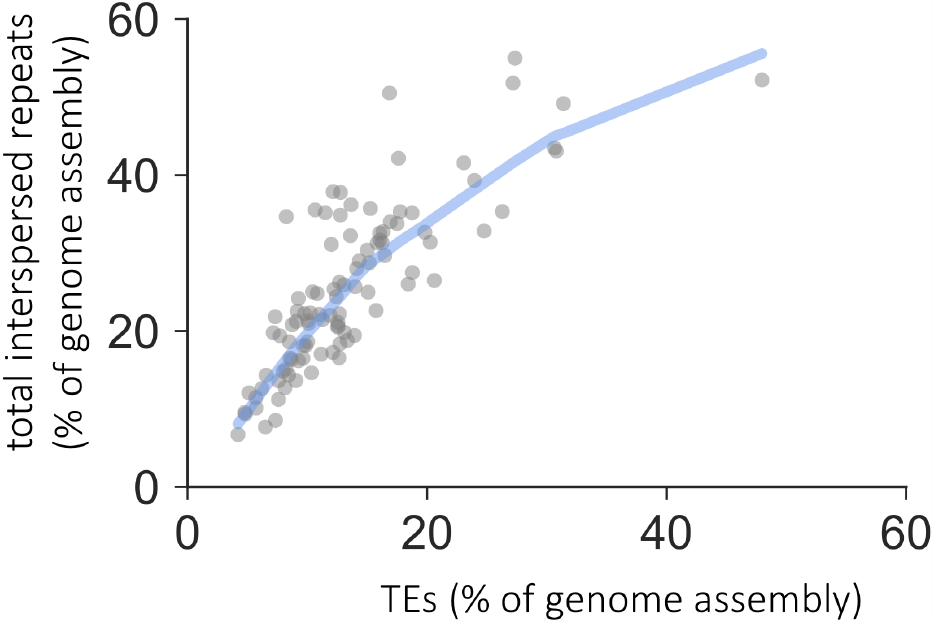
The relationship between the genomic percentage of total interspersed repeats and the percentage of elements classified as TEs.

**Supplementary Figure 3.**
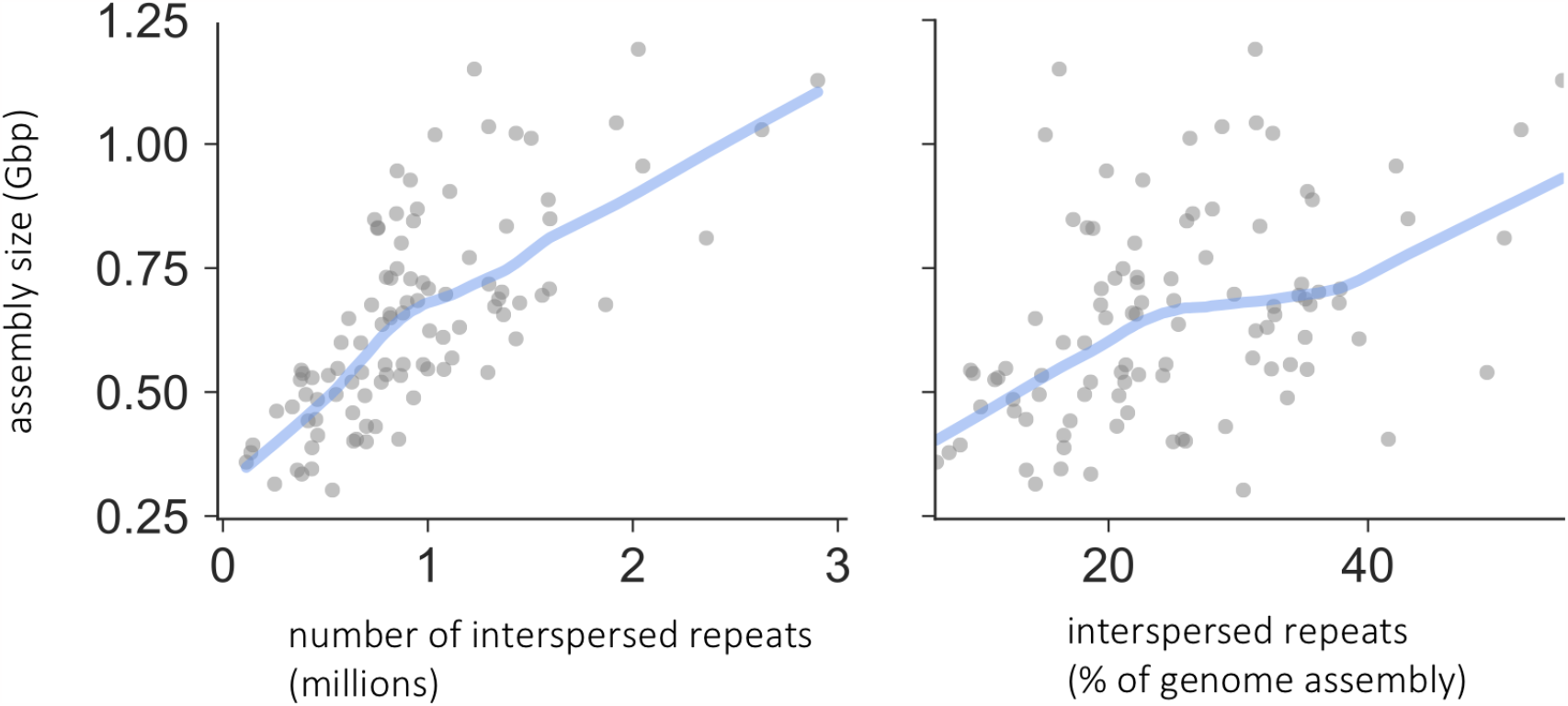
Associations between the number of interspersed repeats (left) and the genomic percentage of interspersed repeats (right) with genome assembly size.

